# A feeding aggregation of Omura‘s whale *Balaenoptera omurai* off Nosy Be, Mozambique Channel

**DOI:** 10.1101/311043

**Authors:** Pierre Laboute, Philippe Borsa

**Affiliations:** Institut de recherche pour le développement (IRD), Anse Vata, BPA5, 98848 Noumea, New Caledonia

## Abstract

A feeding aggregation of Omura’s whales *Balaenoptera omurai* was documented off Nosy Be Island at the northeastern entrance of Mozambique Channel in November 1994. Underwater photographs of live individuals illustrated sub-surface skimming as main feeding behaviour, with small crustaceans, small jellyfish, and other gelatinous micronecton identified as prey. A precise description of the whales’ pigmentation patterns completes previous descriptions from the recent literature.

---

The recently discovered Omura’s whale *Balaenoptera omurai* Wada, Oishi and Yamada 2003 has been reported from, mainly, the tropical waters of the Atlantic, Indian and western-Pacific Oceans (Fig. 1). The osteology of the skull has been described in detail from a number of specimens from the Indian and Pacific oceans^6,7,14,15^. DNA markers have been used to ascertain identification in a number of captured or stranded^9–13,16^ as well as live, Omura’s whales^1,4^. Cerchio and co-authors^4^ have reported regular sightings of Omura’s whales (whose identity was validated by mitochondrial DNA sequence) along the northwestern coast of Madagascar, mainly off the Ampasindava peninsula and also north of Nosy Be island in 2007–2014. We recently exhumated a series of diapositives, taken by one of us (PL), of a then unidentified balaenopterid whale off Nosy Be in 1994. The present note is a brief account of the observations made on these whales, now identified as Omura’s whales in the present report. We describe some of their external morphological features and their pigmentation patterns, and we provide details on their feeding behaviour, thus adding to our knowledge of the external aspect and behaviour of Omura’s whale.

**Figure 1.**
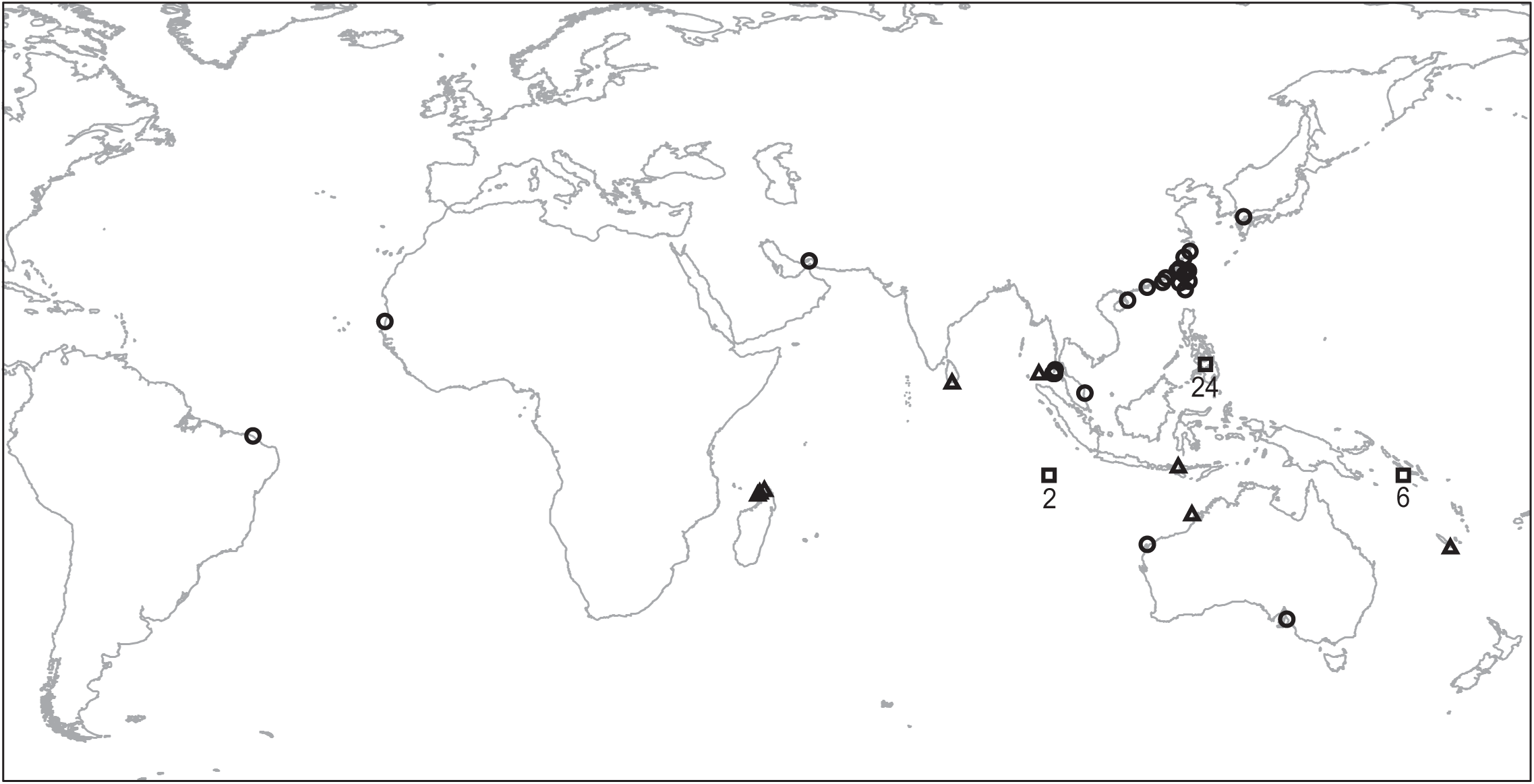
Point-map distribution of Omura’s whale as obtained from sightings^1–5^ symbolized by triangles; strandings^6–13^ (also including three presumably stranded specimens reported from Thailand, and six from Taiwan^14^) symbolized by circles; and captures^6,15^ symbolized by squares, with number of specimens reported.

## Methods

From July 1991 to December 1995, and again in June-August 1998 and June-August 2000 one of us (PL) undertook field work along the reef plateau west of Nosy Be island (northern Mozambique Channel), using a 5-meter outboard motorboat for daily to weekly outings at sea. No survey was conducted during the January and February months. Rorquals of moderately large size, from approximately 8 m to approximately 12 m, were observed on almost every occasion, from June to September every year and occasionally in October and November. Aggregations of whales, with, on 06 November 1994 ca. 10:00 local time, up to a dozen individuals within a radius of less than 300 m, were sighted. The coordinates were 13°26’S 48°05’E. The bottom depth was between 80 m and 40 m, sloping gently towards the shallower reef plateau westwards. On that occasion, the boat was stopped and PL free dived amid the whale aggregation for about one hour to take underwater pictures. Three of the pictures are presented here (Fig. 2) to illustrate the whales’ external morphological features, pigmentation patterns, and feeding behaviour.

**Figure 2.**
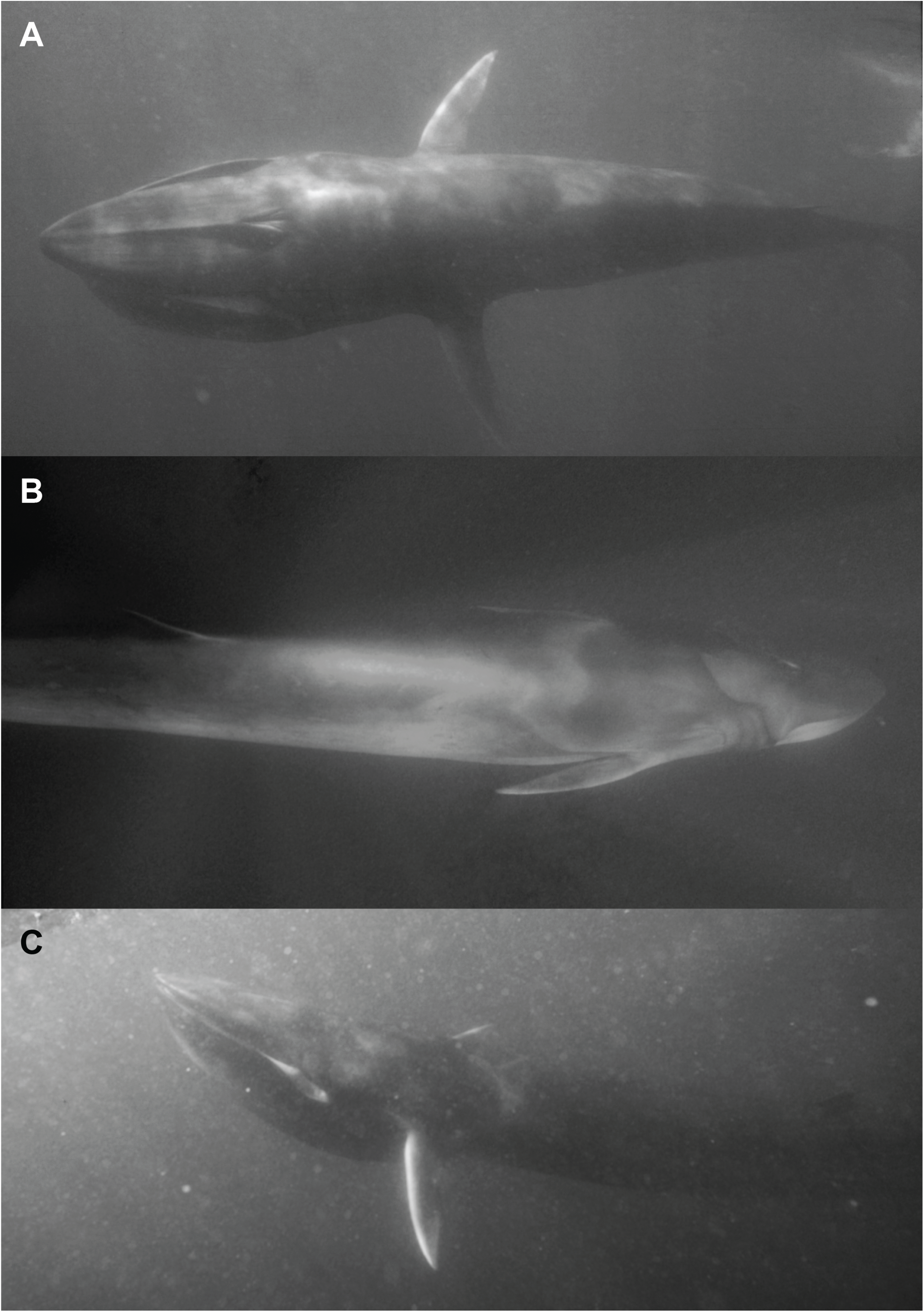
Omura’s whales photographed off Nosy Be (13°26’S 48°05’E), 06 November 1994 by PL. **A**. Skimming posture of Omura’s whale, viewed from above. **B**. View of the right and dorsal sides of an individual. **C**. Backlight view of the left side of another individual, also showing a swarm of gelatinous micronecton under the surface.

## Results and Discussion

Seen from above, the whales’ maxilla had a slightly pointed ogival shape (Fig. 2A). The rostrum bore a single median ridge, and two or three subtle lateral grooves were visible, parallel to the median ridge each side of it (Fig. 2A). The dorsal fin was relatively small, falcate and recurved, its tip pointing backwards (Fig. 2C). The pigmentation of the right mandible was light (Fig. 2B) while that of the left mandible was dark (Fig. 2C). Asymmetry in pigmentation was also visible on the inner part of the lip, which was dark on the right and light on the left (Fig. 2A, C). A series of three parallel, S-shaped dark chevrons connecting the eye and the posterior edge of the mouth to the nuchal region was visible on the right side (Fig. 2B), without matching symmetrical chevrons on the left side (Fig. 2B, C). A lighter-grey, Z-shaped chevron connecting the axil to the anterior part of the back below the shoulders was visible on the right side (Fig. 2B) and a symmetrical, S-shaped chevron was visible on the left side (Fig. 2B, C). The flipper’s anterior edge was light-pigmented, contrasting with the darker pigmentation of the flipper’s dorsal side (Fig. 2A-C).

The whales were observed feeding on a large swarm of zooplankton and micronecton that included small crustaceans, small jellyfish and other gelatinous organisms. The main mode of feeding was by skimming, with the animal gliding at shallow depth (less than 5 m below surface), mouth slightly gaping, allowing the flow of filtered seawater to escape laterally along the gaping rear extremity of the mandible (Fig. 2A, C). Lunge feeding was also observed occasionally, when the rorqual opened its mouth wide and gulped large quantities of gelatinous micronecton. On one occasion, a whale was observed defecating: the animal stood immobile in an upright position below the surface, allowing the plume of faeces to sink.

The whales documented in the present report were identified as Omura’s whales on the basis of their external morphology and pigmentation, including the markedly recurved dorsal fin, the ogival shape of the rostrum, the single median ridge, and the asymmetry in pigmentation patterns^4,6,10^. Pigmentation patterns were very similar between the whales photographed by PL in 1994 (present report) and those photographed in the same area 18–20 years later^4^. In particular, two individuals presented on Cerchio et al.’s figure 3^4^ exhibited a series of three parallel, dark-grey chevrons similar to that photographed on Fig. 2B. Sample monomorphism in the control-region sequences (*N* = 11)^4^ indicates a low genetic diversity for the population of Omura’s whales off northwestern Madagascar, suggesting a low effective population size. This in turn may explain similarities in pigmentation patterns such as those we observed and which at first sight appeared to differ from those of a few individuals from other areas where pigmentation patterns were partly visible (Western Australia, Persian Gulf)^10,11^. However, the skin of the Western Australian specimen had likely undergone post-mortem darkening^10^ and the skin of the Persian Gulf specimen presented cuts, scratches and abrasions on part of its surface (we ascribe these injuries to a collision with a ship), erasing part of the pigmentation^11^. The fact that a similar three-dark chevrons pattern also characterized an individual documented from off Komodo island^1^ and another one from off southern Sri Lanka^5^ suggests that it is, at least, a pattern frequently encountered in Omura’s whales from the Indian-Ocean.

The occurrence of medium-sized rorquals including Omura’s whales off Nosy Be every winter and spring for five consecutive years (present report) and 20 years later again for three consecutive years ^4^ may be either related to seasonal and perhaps year-round suitable trophic conditions, or to suitable conditions for reproduction, or both. Sightings of mother-calf pairs^4^, acoustic records interpreted as courtship vocalizations^4^ and the documentation of feeding behaviour (present report) suggest that the continental shelf waters at the northeastern entrance of Mozambique Channel harbour a permanent or semi-permanent breeding population of Omura’s whale. The interaction of the westward-flowing extension of the South Equatorial current with the topography of the northeastern entrance of the Mozambique Channel generates large anticyclonic eddies^17^. Anticyclonic eddies provoke upwellings which in turn favour high phytoplankton production. Eddies and currents play an important part in the spatial distribution of the chlorophyll in the Mozambique Channel^18^. High chlorophyll concentration associated with upwellings are found along the coasts of Madagascar and Mozambique, notably at the northwestern and southwestern tips of Madagascar^19^. It is possible that the particular topography of the shelf area off northwestern Madagascar favours the accumulation and retention of drifting, fast-growing zooplankton and micronecton including small crustaceans, small jellyfish and other gelatinous animals, which Omura’s whales exploit. Although poorly energetic, this diet may suffice to contribute a part of the energetic requirements of middle-sized rorquals such as Omura’s whales in tropical waters, where individual energetic expenditure is less than in colder waters.

